# Novel repellents for the blood-sucking insects *Rhodnius prolixus* and *Triatoma infestans*, vectors of Chagas disease

**DOI:** 10.1101/789602

**Authors:** Melanie Ramírez, Mario I. Ortiz, Pablo Guerenstein, Jorge Molina

## Abstract

**Background:** Studying the behavioral response of blood-sucking, disease-vector insects to potentially repellent volatile compounds could shed light on the development of new control strategies. Volatiles released by human facial skin microbiota play different roles in the host-seeking behavior of triatomines. We assessed the repellency effect of such compounds of bacterial origin on *Triatoma infestans* and *Rhodnius prolixus*, two important vectors of Chagas disease in Latin America.

**Methods:** Using an exposure device, insects were presented to human odor alone (negative control) and in the presence of three individual tested compounds (2-mercaptoethanol, dimethyl sulfide and 2-phenylethanol, which was only tested in *R. prolixus*) and the gold-standard repellent NN-diethyl-3-methylbenzamide–DEET (positive control). We quantified the time the insects spent in the proximity of the host and performed nonparametric statistical tests to determine if any of the compounds evaluated affected the behavior of the insect.

**Results:** We found volatiles that significantly reduced the time spent in the proximity of the host. These were 2-phenylethanol and 2-mercaptoethanol for *R. prolixus*, and dimethyl sulfide and 2-mercaptoethanol for *T. infestans*. Such an effect was also observed in both species when DEET was presented, although only at the higher doses tested.

**Conclusions:** The new repellents modulated the behavior of two Chagas disease vectors belonging to two different triatomine tribes, and this was achieved using a dose up to three orders of magnitude lower than that needed to evoke the same effect with DEET. Future efforts in understanding deeply the mechanism of action of repellent compounds such as 2-mercaptoethanol, as well as an assessment of their temporal and spatial repellent properties, could lead to the development of novel control strategies for insect vectors refractory to DEET.

## Introduction

Most vectors of human infectious diseases are bloodsucking insects, and therefore, many of those diseases could be potentially eradicated by insect-vector control strategies [1]. For example, it is strongly advised that people living in or visiting regions populated by insects that feed on blood, such as mosquitoes, should protect themselves using insect repellents [2]. Independent of its mechanism of action, the final effect of a repellent is to cause an insect to make oriented movements away from its source. The expected result is to disrupt the host-seeking behavior of the threatening insect [3–5].

Triatomine bugs (Hemiptera: Reduviidae: Triatominae) feed on the blood of vertebrates and are vectors of the protozoan parasite *Trypanosoma cruzi*, the etiological agent of Chagas disease, also known as American trypanosomiasis [6]. The vast majority of the extant 149 species of triatomines are found in Latin American countries, where 68 triatomine species have been found infected with *T. cruzi*, and more than 150 species of domestic and wild mammals have been found to carry the parasite [7,8]. However, few triatomine species are recognized as competent vectors, and only approximately five species are considered very important vectors for humans: *Rhodnius prolixus* Stål, 1859 (inhabiting mainly Colombia and Venezuela), *Triatoma infestans* (Klug, 1834) (inhabiting mainly Peru, Bolivia, Paraguay, and Argentina), *T. dimidiata* (Latreille, 1811) (inhabiting Mexico and Central America), *T. brasiliensis* Neiva, 1911 and *Panstrongylus megistus* (Burmeister, 1835) (both found mainly in Brazil) [8,9]. The infection can occur if, after taking a large blood meal, the insect defecates on the host skin and the feces carrying infective forms of *T. cruzi* enter the blood stream through the wound or any mucous tissue [8]. Since its discovery by Carlos Chagas, until now, controlling vectorial transmission has been the most suitable method to prevent Chagas disease, which affects approximately 7 million people worldwide [10].

Historically, most research on repellents has focused on mosquitoes over other blood-sucking arthropods such as triatomines [4,11–17]. This tendency to focus on mosquito– repellent research is not surprising considering the higher mortality and morbidity due to mosquito-borne diseases compared to Chagas disease [18–20]. For almost six decades, NN-diethyl-3-methylbenzamide, known as DEET, has been the most common mosquito repellent used worldwide [21]. In fact, the effectiveness of DEET against all groups of biting arthropods, triatomines included, has granted it the title of the *gold standard* among repellents [4,5]. However, compared with mosquitoes and other blood-sucking arthropods, triatomines have a lower sensitivity to this repellent [15,22]. Studies with *R. prolixus* and *T. infestans* have revealed that whether the host is present or not, only high doses (i.e., >90%) have a repellent effect, making DEET rather impractical for reducing human-vector contacts [11,23–26]. In addition to these and other related findings in triatomines (i.e., DEET pre-exposure adaptation, DEET repellency in pyrethroid resistant colonies and the effect of nitric oxide on the sensory detection of DEET) [16,27,28], other studies have explored natural repellents such as essential oils, aiming at finding alternatives to DEET and other synthetic repellents [14,17,19,29–32].

A decade of research has shown that volatile organic compounds (VOCs) from human skin and of microbial origin play a role in the behavioral responses of some blood-sucking insects [33]. For example, VOCs produced by skin bacteria are important cues for the malaria vector *Anopheles gambiae* in identifying hosts as human and confer specificity to certain body regions on which mosquitos tend to bite more [33–38]. Moreover, previous studies carried out in our laboratory have demonstrated the role that VOCs released by human facial skin microbiota play in the host-seeking behavior of *R. prolixus* [39–41]. Tabares and collaborators [39] showed, in dual choice olfactometer experiments, that VOCs produced *in vitro* by some skin bacteria (at specific growth phases) had an attractive effect on *R. prolixus*. The authors also reported odor-source avoidance when some other bacteria VOCs were presented, such as those produced by *Citrobacter koseri* (Enterobacteriaceae). Insects consistently chose the negative control (i.e., culture medium without bacteria) over the culture medium with bacteria VOCs. These two findings, the attractive and avoidance behavioral effects, contrasted with those from other bacterial VOCs to which *R. prolixus* did not respond at all [39].

Therefore, the behavioral response of triatomines to the mix of VOCs produced by the skin microbiota seems to be very complex [39]. Moreover, the role of individual bacterial volatiles from mixtures in evoking avoidance is still unknown, and their potential use as repellents deserves further investigation. In this study, we asked whether individual VOCs released by cultures of *C. koseri*, which evokes avoidance, could affect the behavior of kissing bugs in the proximity of a human host exhibiting, for example, a repellent effect.

Furthermore, we investigated whether this potential effect could be equivalent to that evoked by the well-known repellent DEET. Thus, using an exposure device, we assessed in *R. prolixus* and *T. infestans* the repellency effect of three compounds whose chemical structure is similar to that of compounds identified from cultures of *C. koseri* [36]: 2-mercaptoethanol, 2-phenylethanol and dimethyl sulfide. We compared the repellency effectiveness of these compounds at different doses with that obtained with DEET.

## Methods

### Insects

Adults of *R. prolixus* and third-instar nymphs of *T. infestans* from our laboratory colonies were used. The *R. prolixus* colony originated from wild populations from San Juan de Arama, Meta Department (Northeast of Colombia), and has been maintained at the Centro de Investigaciones en Microbiología y Parasitología Tropical–CIMPAT in Universidad de los Andes (Bogotá, Colombia), while the *T. infestans* colony originated from wild populations from Chaco province (Northeast of Argentina; provided by the Servicio Nacional de Chagas of Argentina), and has been maintained at the Centro de Investigacion Cientifica y de Transferencia Tecnologica a la Produccion (CICyTTP, Diamante, Argentina). Insects were fed on hens every two weeks and maintained under an artificial 12:12 (L:D) illumination regime at controlled temperature and humidity (27± 2°C, 75 ± 10% RH).

For experiments, insects were separated from the colony after molting and starved for at least 20 days for *R. prolixus* and 30 days for *T. infestans*. Experiments were video recorded (using a DCR-SR 200 camera (Sony Corp., Japan) or an A1633 iPhone camera (Apple Inc., USA)) and performed during the early scotophase at 24.5 ± 0.5 °C in a dark (or red-light illuminated) room. Experiments with *R. prolixus* were performed at CIMPAT, Universidad de los Andes, and experiments with *T. infestans* were carried out at Laboratorio de Estudio de la Biología de Insectos – LEBI, CICyTTP-CONICET. Insects were tested individually and used only once.

### Repellency tests

To test the individual effect of compounds produced *in vitro* by bacteria previously isolated from human facial skin, an exposure device modified from Zermoglio and collaborators [11] was used. In brief, a polystyrene tube was divided into three zones: host, intermediate and refuge zones. The host stimulus consisted on a human forearm. Insects were placed in the refuge zone, and after a five-minute adaptation time, the experiment started with the opening of a gate, allowing the insect to freely move from the refuge to the other two zones. Insects attracted by the stimuli from the host walked to the host zone, while a mesh prevented them from biting the forearm. Experiments lasted five minutes. The exposure device allowed us to quantify the time the insect spent near the host in the presence or absence of the compounds tested.

Ten insects per treatment were used; these were randomly assigned to each treatment. Treatments for experiments with *R. prolixus* consisted of 2-mercaptoethanol (0.0015625%, 0.003125%, 0.00625%, 0.0125%, 0.025%, 0.05% and 0.1%), dimethyl sulfide (0.00625%, 0.0125%, 0.025%, 0.05% and 0.1%), 2-phenylethanol (0.025%, 0.05%, 0.1% and 0.2%), and DEET (10%, 50%, 90%). Treatments for experiments with *T. infestans* consisted of 2-mercaptoethanol (0.00625%, 0.025%, 0.1% and 1%), dimethyl sulfide (0.1% and 1%), and DEET (90%). The tested compounds were ≥99% pure (Merck, Darmstadt, Germany), while DEET was >97% pure (Sigma-Aldrich, Darmstadt, Germany). Dimethyl sulfide and 2-mercaptoethanol solutions were made in distilled water, while ethanol was the solvent for 2-phenylethanol and DEET. We performed frequent negative-control tests: host stimuli without any test compound (“host alone” see below) and host stimuli plus just ethanol (“host plus ethanol”, see below). Test-odor stimuli consisted of a 10 μl solution (or just solvent for the controls) loaded onto a filter paper strip (1.0 x 3.0 cm). In the case of DEET, 10 μl or 50 μl solutions (where indicated) were used. The paper strip with the test solution or solvent control was carefully placed in the space between the host’s forearm and the mesh in the tube. Neither the host’s skin nor the insects were in direct contact with the compounds tested.

### Data analysis and statistics

We carried out nonparametric statistical tests to determine whether the compounds influenced the time that the insect spent in the host proximity. Prism software (GraphPad, 7.0a) was used to perform Kruskal-Wallis and Dunn’s multiple comparison tests (*p* < 0.05) within each treatment group.

## Results

In this study, we assessed the repellency of VOCs released by the skin bacterium *C. koseri* on *R. prolixus* and *T. infestans*. For this, 240 starved adult *R. prolixus* and 90 starved *T. infestans* nymphs were assayed in an exposure device.

In the absence of test compounds (negative controls), *R. prolixus* spent 241 s (“host alone” *a*), 148.5 s (“host alone” *b*), and 255.5 s (“host alone” *c*) in the host zone out of 300 s of experimental time (median values) (Fig. 1, white boxes). In the case of *T. infestans*, insects spent 177 s (“host alone” *d*) (median value) in the host zone out of 300 s of experimental time (Fig. 2, white boxes).

**Figure 1.**
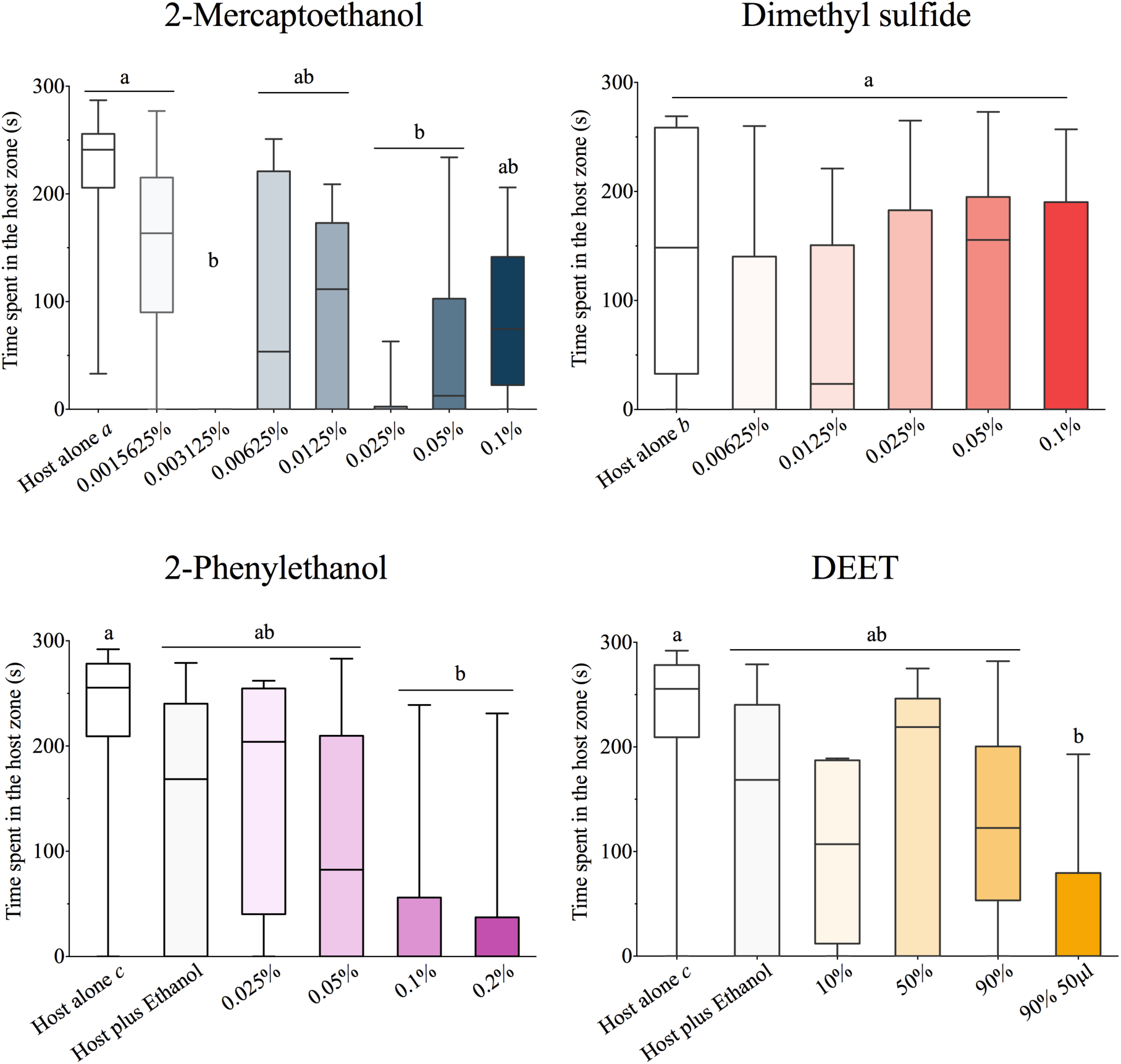
Box plots showing the effect of different doses of the test compounds on the time that *R. prolixus* spent in the proximity of a vertebrate host when the insects were exposed to 2-mercaptoethanol, dimethyl sulfide, 2-phenylethanol, and DEET (median, maximum and minimum values are shown). Letters denote significant differences among treatments according to Dunn’s multiple comparison test (p<0.05). *Host alone a, b* and *c* are repetitions of a negative control consisting of exposure to the forearm of the host in the absence of any test compound.

**Figure 2.**
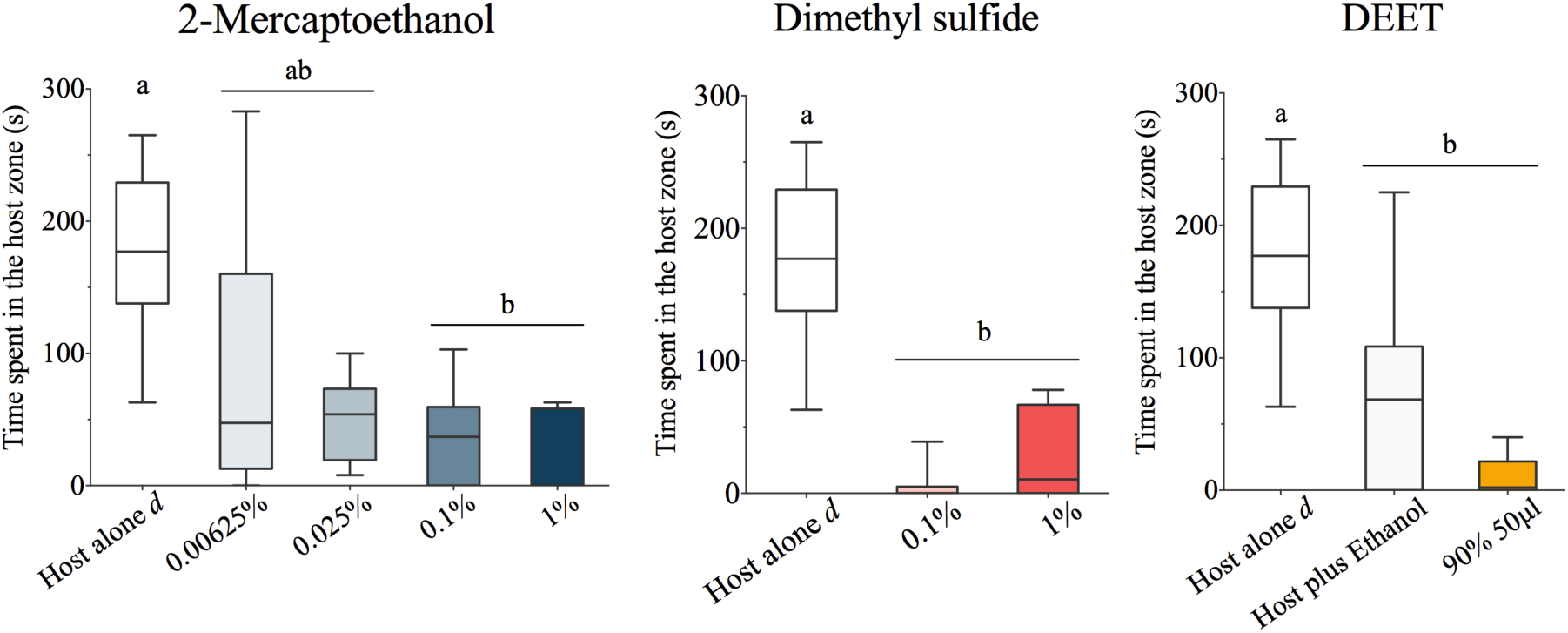
Box plots showing the effect of different doses of the test compounds on the time that *T. infestans* spent in the proximity of a vertebrate host when the insects were exposed to 2-mercaptoethanol, dimethyl sulfide, and DEET (median, maximum and minimum values are shown). Letters denote significant differences among treatments according to Dunn’s multiple comparison test (p<0.05). *Host alone d* refers to a negative control consisting of exposure to the forearm of the host in the absence of any test compound.

However, when certain doses of 2-mercaptoethanol, 2-phenylethanol or DEET were added, the time that adult *R. prolixus* spent in the host zone was significantly lower (Kruskal-Wallis test, *p* <0.0001, *p* = 0.0007, *p* = 0.0037, respectively) (Fig. 1). Likewise, certain doses of 2-mercaptoethanol and DEET considerably reduced the time that *T. infestans* nymphs stayed near the host (Kruskal-Wallis test, *p* = 0.0002 and *p* = 0.0002, respectively) (Fig. 2). It should be noted that Dunn’s Multiple Comparison tests showed no differences between the times for treatments in which the compounds were dissolved in ethanol and those for the control “host plus ethanol”. However, a significant difference was found when the former times were compared with those for the host alone (Table 1).

**Table 1.**
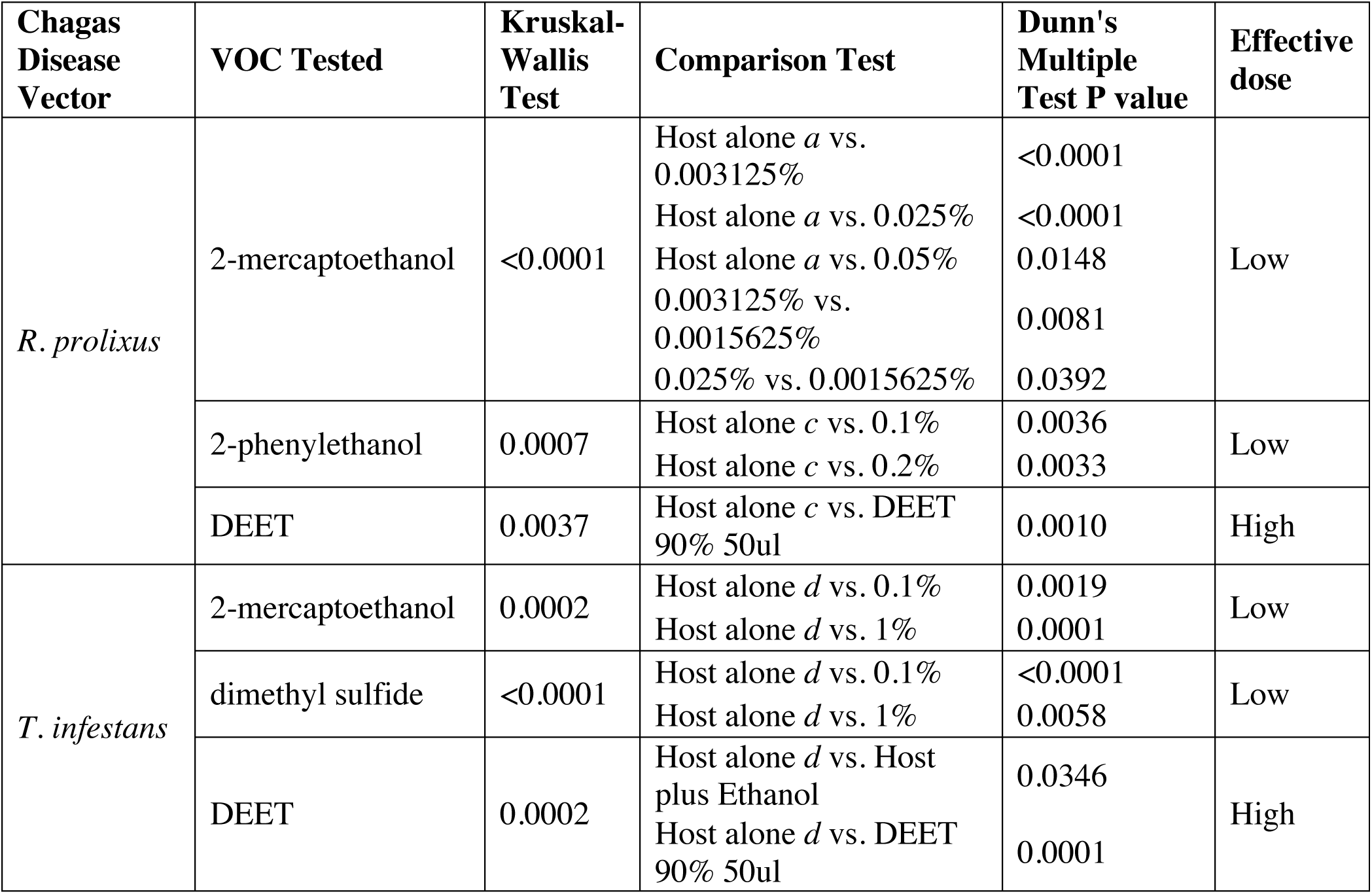
Summary of the multiple comparisons tests that resulted in statistically significant differences (p<0.05), showing treatments that reduced the time the insects spent in the host zone with respect to a negative control.

| Chagas Disease Vector | VOC Tested | Kruskal-Wallis Test | Comparison Test | Dunn's Multiple Test P value | Effective dose |
| --- | --- | --- | --- | --- | --- |
| *R. prolixus* | 2-mercaptoethanol | <0.0001 | Host alone *a* vs. 0.003125%<br>Host alone *a* vs. 0.025%<br>Host alone *a* vs. 0.05%<br>0.003125% vs. 0.0015625%<br>0.025% vs. 0.0015625% | <0.0001<br><0.0001<br>0.0148<br>0.0081<br>0.0392 | Low |
|  | 2-phenylethanol | 0.0007 | Host alone *c* vs. 0.1%<br>Host alone *c* vs. 0.2% | 0.0036<br>0.0033 | Low |
|  | DEET | 0.0037 | Host alone *c* vs. DEET 90% 50ul | 0.0010 | High |
| *T. infestans* | 2-mercaptoethanol | 0.0002 | Host alone *d* vs. 0.1%<br>Host alone *d* vs. 1% | 0.0019<br>0.0001 | Low |
|  | dimethyl sulfide | <0.0001 | Host alone *d* vs. 0.1%<br>Host alone *d* vs. 1% | <0.0001<br>0.0058 | Low |
|  | DEET | 0.0002 | Host alone *d* vs. Host plus Ethanol<br>Host alone *d* vs. DEET 90% 50ul | 0.0346<br>0.0001 | High |

The time spent by *R. prolixus* near the host did not differ statistically from the negative control when dimethyl sulfide was tested (Kruskal-Wallis test, *p* = 0.1414). In contrast, dimethyl sulfide did reduce the time that the *T. infestans* nymphs spent near the forearm (Kruskal-Wallis test, *p* <0.0001). A summary of the statistically significant results of the multiple comparisons tests is shown in Table 1.

## Discussion

Our results provide evidence that some VOCs released by the opportunistic skin bacterium *Citrobacter koseri* interfere with the host-seeking behavior of *R. prolixus* and *T. infestans*, two important vectors of Chagas disease. In negative control tests where just a host is presented, *R. prolixus* adults and *T. infestans* nymphs move their antennae in a triangulation fashion [42,43], and in just a few seconds, walk towards the host, extend their proboscis and insistently try to bite the forearm. However, when the compounds tested are added to the stimuli of the host, the behavior of the bugs changes; the time spent near the human host is considerably reduced (see Results), and the frequency of biting attempts is lower (data not shown). Moreover, our observations show that the reduction in the time spent in the proximity of the host is because there is an augmentation of the latency time (i.e., insects are delayed in making the decision to move forward) and they spend a very short time in the host zone (i.e., insects going in and out of the host zone). An additional movie file shows that both species rapidly walk away from the stimulus source after approaching it [see Additional file 1]. Suggesting a clear repellent effect on triatomines working against potential attractive stimuli like thermo and chemoreception mediated by host VOCs. The methodology used in this work (based on that by Zermoglio and collaborators [11]) suggested a fast and direct way to test the effect of candidate VOCs on the repellency of triatomines when the VOCs were applied near a vertebrate host.

The fact that *R. prolixus* is attracted by some VOCs released by some common bacteria of the human face skin, as Tabares and collaborators [39] showed, could be related to the close vertebrate-vector coevolutive history. However, the response of the bugs to the VOCs produced by *C. koseri* could make sense if the natural occurrence of the bacterium is considered: *C. koseri* is a gram-negative bacillus of the Enterobacteriaceae family, found in animal intestines, soils, water, sewage and contaminated food, and widely recognized for causing devastating meningitis in neonates and severe infections in immunosuppressed patients [44]. As this bacterium is not part of the healthy human skin microbiota (human skin isolations where this bacillus was found are commonly from sick patients, see [44,45]), blood-sucking insects such as triatomines would barely contact the volatile products of the bacterium. Moreover, it could signal an unhealthy individual to the bugs.

Interestingly, it is not new that the VOC signature of the genus *Citrobacter* influences the chemotactic orientation behavior of blood-seeking insects. Ponnusamy and collaborators found that VOCs released by *Citrobacter freundii* were attractive to gravid females of *Aedes (Stegomyia) aegypti* and *Ae. (Stegomyia) albopictus* [46]; both mosquitoes are well recognized as vectors of important arboviruses [47]. It was also suggested that *Citrobacter* VOCs, in synergy with other compounds present in water, give mosquito information about the quality of the oviposition sites [46]. In the bloodsucking stable fly *Stomoxys calcitrans*, Romero and collaborators showed that *C. freundii* was a strong cue inducing oviposition in soil [48]. Therefore, VOCs released by *Citrobacter sp*. appear to be an interesting semiochemical source, mediating interactions with biotic (e.g., animal and human hosts) and abiotic (e.g., water and soil) factors, which is crucial for insects of medical importance [49–52].

The VOC mix released by *Citrobacter* sp. can be described as having a strong, fetid and putrid odor. Many species among the genus are cataloged within the malodor-generating bacteria group, in part because of their participation in decomposition processes [53-55]. The compounds methanethiol and dimethyl disulfide, identified as VOCs released by *C. koseri* [39], and the two VOCs used in our study, 2-mercaptoethanol and dimethyl sulfide, are sulfur-containing compounds with a strong smell. Sulfur compounds are neurotoxic and lethal to some insects and are proposed as a new control alternative to agricultural pests [56,57]. However, in addition to the repellency effect, we did not identify any symptoms of intoxication (i.e., insects with abnormal rest positions, paralysis in the legs or death. [15]) due to sulfur compounds in our experiments, perhaps because of the low doses tested; nevertheless, the toxicity of these sulfur compounds to animals and humans should be reviewed carefully for future applications.

Both sulfur compounds, together with 2-phenylethanol, are also known and used as VOC markers of human and animal wastes [58,59]. They are also involved in the decomposition of mammal and bird tissues [60,61], a scenario that is not very attractive to triatomine insects. It is interesting to note that 2-phenylethanol is also produced by the Brindley’s gland of *T. infestans*, a gland involved in the alarm pheromone production of the adult [62–65]. However, this compound has not been reported as part of the alarm pheromone of *R. prolixus* [63]. In this study, we showed that 2-phenylethanol has a repellent effect on *R. prolixus*. Likewise, in *An. gambiae*, this compound was reported as a spatial repellent candidate that inhibits attraction [66,67]. The effect that this compound could have on the behavior of *T. infestans* needs to be further assessed. It should be noted that in this work, the time spent in the host zone when presenting 2-phenylethanol was significantly lower than that of the “host alone” negative control but not different from the “host plus ethanol” negative control. Additionally, there were no significant differences between the two negative controls. This suggests that the repellent effect of 2-phenylethanol is evident only when presented together with ethanol, possibly due to a synergistic effect between the solvent and the test compound.

In this work, DEET was used as a positive control. As our results show, the repellency effect of DEET for *R. prolixus* may be the result of a synergy between the solvent and DEET, as in the case of 2-phenylethanol. Such a repellency effect of DEET (plus ethanol) was only achieved at the highest dose tested (i.e., 90%–50μl). In contrast, 2-phenylethanol (for *R. prolixus*), dimethyl sulfide (for *T. infestans*) and 2-mercaptoethanol (for both species) showed a repellent effect at doses two to three orders of magnitude lower than the effective dose of DEET (i.e., 0.003125% – 0.1%). Efficiency at low doses is one of the key characteristics that is required for a good, new repellent [21]. The need to employ high concentrations of DEET to achieve repellency has limited its application in disrupting triatomine-human contacts, as several studies have already shown [11,23–26]. Although its use is deemed safe, DEET has some disadvantages: it needs to be constantly reapplied, it has a short range of action due to its low volatility and can melt plastics and vinyl [4,21]. Even more important, the people who truly need it usually cannot afford it [4].

The question of why triatomines are almost refractory to the gold standard DEET is still open. One hypothesis concerning the repellent effect of DEET is that it mimics a defensive compound of plants, methyl jasmonate, explaining why in insects with a large-plant evolutive history, such as mosquitoes, it is still effective [4,68]. Although some triatomine species such as *Rhodnius prolixus* have a close relationship with palm tree niches [71], resting within such plants although not feeding on them, molecules as DEET may not be directly related to the triatomine evolutive history as it is with mosquitoes (i.e., early ancestors of the Triatominae subfamily were predators, unlike plant-feeder mosquito ancestors). In fact, triatomines are obligate hematophagous, and many species have nearly zero contact with plants [26,69–71]. Despite the advances in research on repellency in mosquitoes, where DEET is considered the gold standard, finding efficient repellents for triatomines still represents a challenge.

## Conclusions

As far as we know, this is the first study in triatomines that assesses the repellent effect of individual volatiles of microbial origin from a human host. We showed that vectors of two different tribes (Rhodniini and Triatomini), with epidemiological importance in Chagas disease transmission, are repelled by very low doses of the sulfur compound 2-mercaptoethanol. Future studies should be directed to understand deeply its mechanism of action in triatomines and to assess its possible use as a repellent (although not applied directly onto the skin) or within a push-pull control strategy.

## Acknowledgements

MR is grateful to Colciencias and Facultad de Ciencias–Universidad de los Andes for funding this project and fellowship (Convocatoria Nacional para estudios de Doctorados No. 567, and Proyecto Semilla 2018 for Candidate PhD. Students). JM is also grateful to Colciencias (funding project 759-2013). PG acknowledges Agencia Nacional de Promoción Científica y Tecnológica (ANPCyT, Argentina) for funding through grant PICT 2015, N° 3260.

## Additional file 1.mp4

Video recording showing the repellency effect of tested compounds on *R. prolixus* and *T. infestans*. The time in the host zone was diminished either by augmenting the time to get in the host zone or by moving in and out of the host zone.

